# Intrinsic Error Correction in Protein Allostery: Quantifying Noise Suppression via Spanning Tree Statistics

**DOI:** 10.64898/2025.12.22.696126

**Authors:** Burak Erman

**Affiliations:** Chemical and Biological Engineering, Koc University, Istanbul, Turkey

## Abstract

Allosteric regulation in proteins arises from collective dynamics distributed over networks of residue contacts, but how multiple communication pathways contribute to signal transmission and noise suppression remains unclear. Here we develop a spanning-tree–based framework to quantify allosteric communication as an ensemble of interacting pathways in protein contact networks.

We introduce a dynamic distance measure linking local perturbations of residue interactions to global changes in network entropy, establishing a local-to-global scaling between local dynamics and global sensitivity. Using spanning-tree calculus, we derive exact probabilities for all simple paths connecting pre-specified functional residue pairs. This enables a comparison between an approximate description based on uniform path usage and a topology-aware description in which path probabilities are determined by the Burton-Pemantle theorem and reflect network dependencies.

From these path ensembles, we define corresponding signal-to-noise ratios and quantify how pathway multiplicity and statistical weighting shape noise suppression. Applied to KRAS and to 20 additional allosteric proteins spanning diverse functional classes, the analysis shows large variability in path usage, entropy reduction, and signal-to-noise enhancement, while consistently demonstrating that topology-aware weighting concentrates signal transmission onto dominant short pathways. This suggests that the robustness of allosteric signaling is a fundamental emergent property of protein contact topology.

These results provide a quantitative framework linking protein structure, dynamics, and information flow, and show that robustness in allosteric communication, manifested as noise suppression through pathway redundancy, can be interpreted as an intrinsic error-correction or noise averaging mechanism arising from network topology.

## I. Introduction

Living systems maintain their organization by processing, transmitting, and correcting information under constant physical and chemical noise. From genetic replication to neural signaling, every level of biological activity relies on mechanisms that preserve informational fidelity while allowing adaptive flexibility. At the molecular scale, this challenge is exemplified by proteins, which must coordinate distant sites through dense networks of interacting residues while operating under strong thermal noise.

Proteins are not rigid structures but dynamic networks that coordinate fluctuations across widely separated sites. Through these correlated motions, they convert local perturbations‒such as ligand binding, mutation, or environmental change‒into global functional responses [1-5]. Here, “information” is understood operationally as the transmission of such perturbations through collective protein dynamics, rather than as an explicit biochemical or coding variable. This ability to sense, propagate, and regulate signals suggests that proteins act as molecular information-processing systems. Understanding how these capabilities emerge from collective dynamics is a central question in the physics of life.

These ideas have been widely appreciated, and have been discussed in Ref. [6]. A growing body of work [7-10] has provided important qualitative insight into allosteric communication, robustness, and noise filtering in proteins. Building on this foundation, we develop a quantitative framework that makes these concepts explicit and computable. This work highlights that despite recognition of intramolecular protein signaling and communication [11-21], the rigorous quantification of error correction in proteins requires novel theoretical and computational tools beyond traditional biochemical methods.

Computational approaches, such as elastic network models and structure-based molecular dynamics on simplified energy landscapes, have provided key insights into allosteric transitions and dynamic networks [22]. Here we address this challenge by applying spanning tree statistics [23,24] to the Gaussian Network Model (GNM) [25], which captures the collective fluctuation modes encoded by protein contact topology. In the Theory section, we formalize this picture using the matrix‒tree theorem, which provides a principled way to enumerate and weight the ensemble of possible communication pathways between residues. Although spanning trees have previously been used in protein network analysis, most commonly in the form of minimum spanning trees that identify a single optimized backbone [26], and more recently to characterize information transfer [27], no framework exists for quantifying noise suppression through pathway multiplicity. Such quantification requires: (i) computing the complete statistical ensemble of pathways, (ii) assigning exact probability weights to each route, and (iii) deriving signal-to-noise ratios from this weighted ensemble—precisely what our spanning-tree framework provides.

We introduce a dynamic distance measure that captures how strongly the motions of two residues are coupled through the network of collective fluctuations. Conceptually, this measure can be interpreted as a resistance to information transfer between residues i and j, analogous to electrical resistance in a circuit [28,29]. This quantity, denoted *R*_*ij*_, is defined formally in Eqs. (3) and (4).

A central implication of this framework is that allosteric communication does not proceed along a single dominant pathway. Instead, perturbations introduced at one site are transmitted through ensembles of alternative routes formed by the dense contact topology of the protein [6,21,30-33]. These parallel pathways allow the same signal to reach distant residues through multiple dynamically independent channels.

When a signal propagates along multiple pathways, each channel accumulates thermal and dynamical noise independently.

Upon convergence at the receiving residue, noise contributions from different pathways add incoherently, whereas the signal adds coherently through averaging. This separation leads to an enhanced effective signal-to-noise ratio, yielding an intrinsic, topology-driven mechanism of error correction that does not rely on biochemical reaction cycles. We use the term error correction in the information-theoretic sense of reducing error probability through redundancy and noise averaging; no active identification of individual errors is implied.

Using this framework, we identify several general features of intramolecular information transfer. Communication between residues is mediated by ensembles of parallel pathways rather than single dominant routes. Intramolecular communication is heterogeneous, in the sense that edges differ in their probability of occurring in communication paths. Edges with high path probability are frequently used by these pathways and are termed bottleneck edges, whereas edges with low path probability are used less frequently and provide alternative routes for communication, which we term redundant edges. Residues incident on bottleneck and redundant edges are correspondingly referred to as bottleneck and redundant residues. The formal relationship between dynamic distance and path probability is established in Appendix B.

This perspective suggests a formal analogy to engineered communication systems: allosteric perturbations can be viewed as signals transmitted through noisy channels, where the protein’s contact network topology determines channel characteristics. Just as engineered systems use redundancy and error correction to ensure reliable communication, proteins may exploit pathway multiplicity to suppress noise and maintain signaling fidelity. Here we make this analogy quantitative by calculating channel probabilities and signal-to-noise ratios for allosteric communication networks.

We apply this framework to KRAS—a central signaling switch with well-characterized long-range allostery—and extend the analysis to a benchmark set of 20 proteins with experimentally characterized allosteric regulation spanning diverse functional classes. This reveals both universal principles and system-specific variation in how proteins balance pathway multiplicity against information concentration.

Across this benchmark, we observe substantial quantitative diversity. In some proteins, communication pathways are distributed relatively uniformly, leading to high effective signal-to-noise ratios through extensive pathway multiplicity. In others, information transfer is concentrated along a small number of dominant routes, resulting in lower pathway entropy and reduced noise averaging. The present framework provides a unified statistical description of this spectrum of behaviors without invoking system-specific mechanisms.

More broadly, this work reframes protein allostery as a problem of statistical information transfer in a noisy network, placing concepts from network theory and statistical physics on a quantitative footing in molecular biology. In the Theory section that follows, we formalize these ideas by linking collective dynamics, spanning-tree statistics, and information transfer within a unified mathematical framework.

## II. Theoretical Framework

We now formalize the connection between protein contact topology and information transfer capacity. Starting from the Gaussian Network Model and dynamic distance, we use spanning tree statistics to quantify pathway multiplicity and its role in error correction. Detailed derivations appear in the Appendices.

### A. Dynamic distance and local-to-global scaling in the Gaussian Network Model

To quantify how proteins transmit and correct information through their internal dynamics, we formulate the Gaussian Network Model (GNM) directly in the language of graph theory. A protein is represented as an undirected graph G=(V,E), where each residue *i* corresponds to a vertex *i* ∈*V*, and an edge of weight *w*_*ij*_ with (*i, j*)∈ *E* is present if residues i and j form a native contact of (typically defined via a cutoff distance in the crystal structure). The contact topology is encoded in the adjacency matrix *A* with

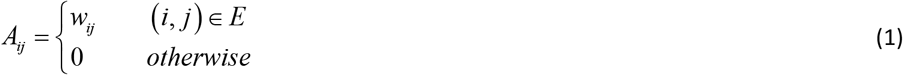

The elastic connectivity of the network is captured by the (combinatorial) Laplacian matrix

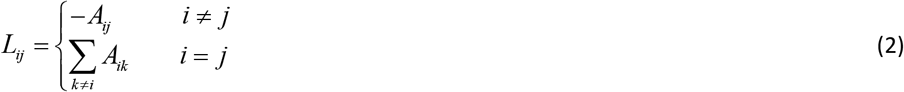

so that *L* = *D* − *A*, with *D* the diagonal degree matrix.

Equation 2 is the form used throughout the paper and the Appendices. In the GNM, *w*_*ij*_ are all taken as unity and the Laplacian serves as the stiffness matrix governing the harmonic fluctuations around the native state.

Because the Laplacian *L* is singular (its nullspace corresponds to rigid-body motions), the relevant dynamics are described by the Moore–Penrose pseudoinverse *L*^+^ . For clarity of notation, we define the pseudoinverse of the graph Laplacian as *K* = *L*^+^ through the paper and appendices, which serves as the fundamental kernel for our analysis. The dynamic distance *R*_*ij*_ between residues i and j is defined as

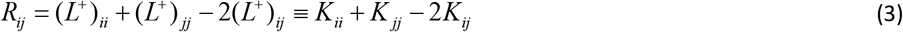

The cross-correlation of fluctuations between residues i and j is proportional to *K*_*ij*_, and within the GNM framework with thermal equilibrium, the pseudoinverse elements directly relate to mean-squared fluctuations through the equipartition theorem [34]

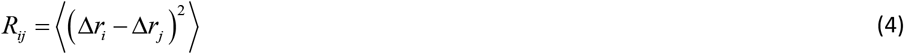

where, the angular brackets denote time average and *r*_*i*_ is the position vector of residue i and Δ*r*_*i*_ is the instantaneous fluctuation from the mean. This identification enables a precise, graph-theoretic decomposition of protein dynamics into contributions from all possible communication pathways. In Appendix B, the Matrix Tree Theorem [24] is used to obtain the dynamic distance as a weighted sum over all spanning trees of the protein contact graph, and thus as a sum over all distinct paths connecting i and j. Thus, each spanning tree contributes a unique path, providing a natural mathematical definition of pathway multiplicity.

As introduced in the Introduction, we classify edges according to their participation in communication pathways as bottleneck or redundant, and refer to residues incident on these edges as bottleneck or redundant residues. We now formalize this classification using spanning-tree statistics.

From Eq. 4 we derive the mean dynamic distance index, *R*_*i*_, of residue i as

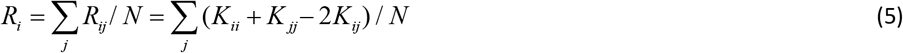

where *N* is the total number of residues represented by their alpha carbons throughout the paper. *R*_*i*_ quantifies the mean dynamic distance of residue i to all other residues in the protein. As shown below using spanning-tree statistics, residues with small *R*_*i*_ are typically incident on edges that are bypassed by many alternative communication paths, whereas residues with large *R*_*i*_ are incident on bottleneck edges through which a large fraction of paths must pass.

In order to understand local and global coupling [35] and separate local effects from global, we define the local versions, *R*_*i,edge*_ as

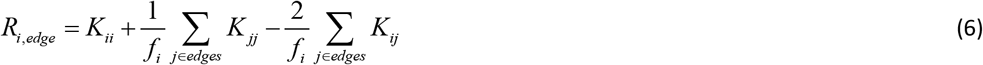

where, *f*_*i*_ is the number of edges incident on residue i. While *R*_*i,edge*_ is a measure of local interactions, *R*_*i*_ carries information on global interactions. In the interest of estimating how much edges contribute to the global, we evaluated *R*_*i*_ and *R*_*i,edge*_ using Eqs. 5 and 6 for 200 proteins and obtained a simple linear scaling equation as

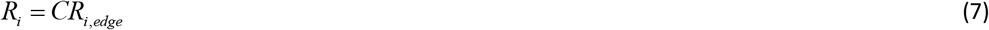

where *C* ≈ 1.93 as may be deduced from the lower right panel of Figure 2.

**Figure 1.**
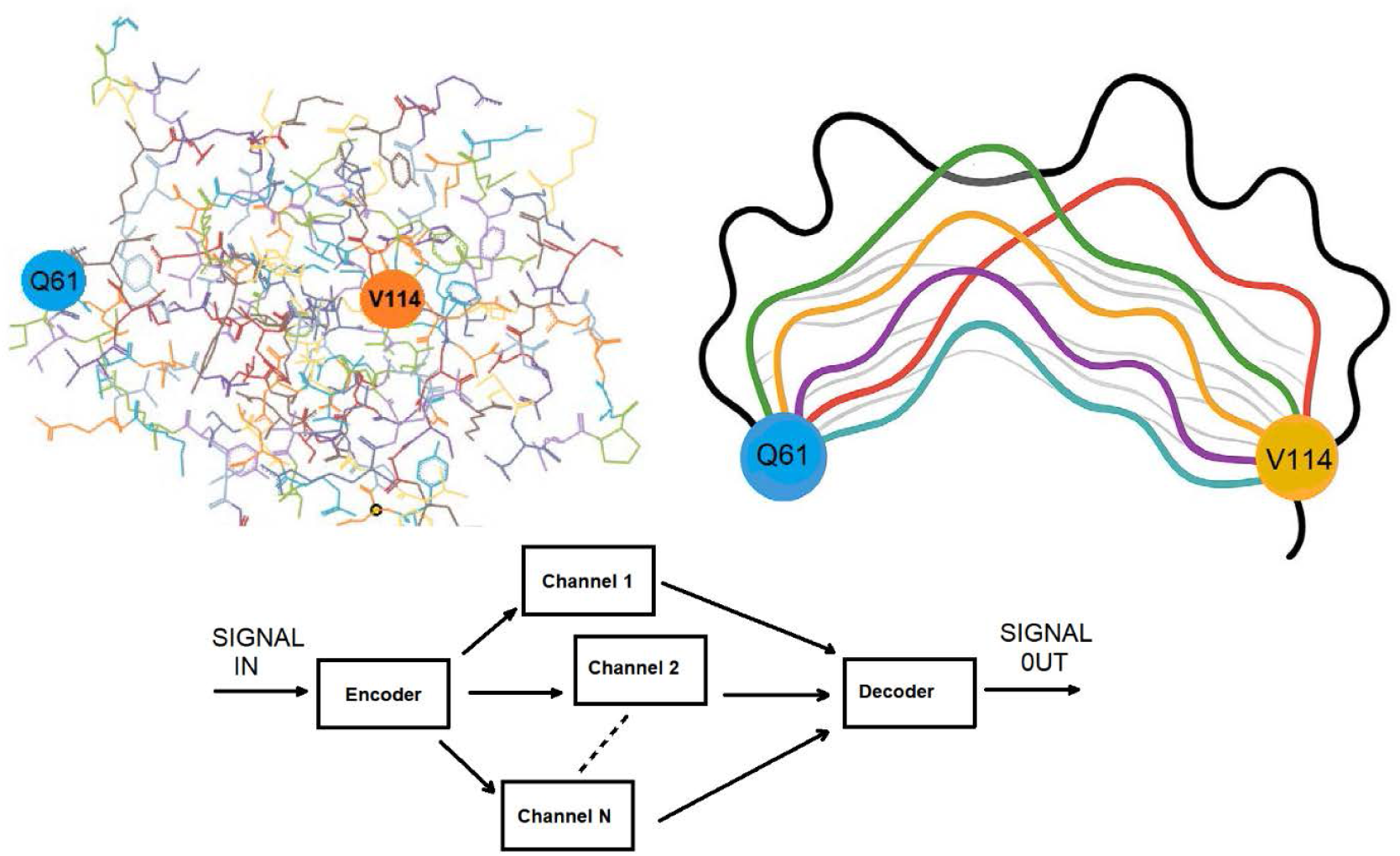
Schematic illustration of information transfer between a redundant residue (V114) and a bottleneck residue (Q61) in KRAS (PDB: 6god). *Upper left:* Wire representation of residues V114 and Q61 showing their structural context within KRAS. V114 is identified as a redundant residue, while Q61 acts as a bottleneck, as detailed in the Results section. Several illustrative information-transfer channels are sketched between the two residues (upper right), with distinct colors indicating channels of different path probabilities; thin, pale lines represent low-probability, low-information, routes. *Lower panel* : Conceptual diagram of the information-transfer process. A sinusoidal input signal is applied at Q61 and is encoded or propagates through by duplication into multiple parallel channels (Channel 1, 2, …, N). Each channel is subject to independent uncorrelated noise. At the receiver side (V114), the signals from all channels are averaged by the decoder, illustrating how multiplicity enhances robustness and reduces the effective noise in information arriving at the bottleneck residue.

**Figure 2.**
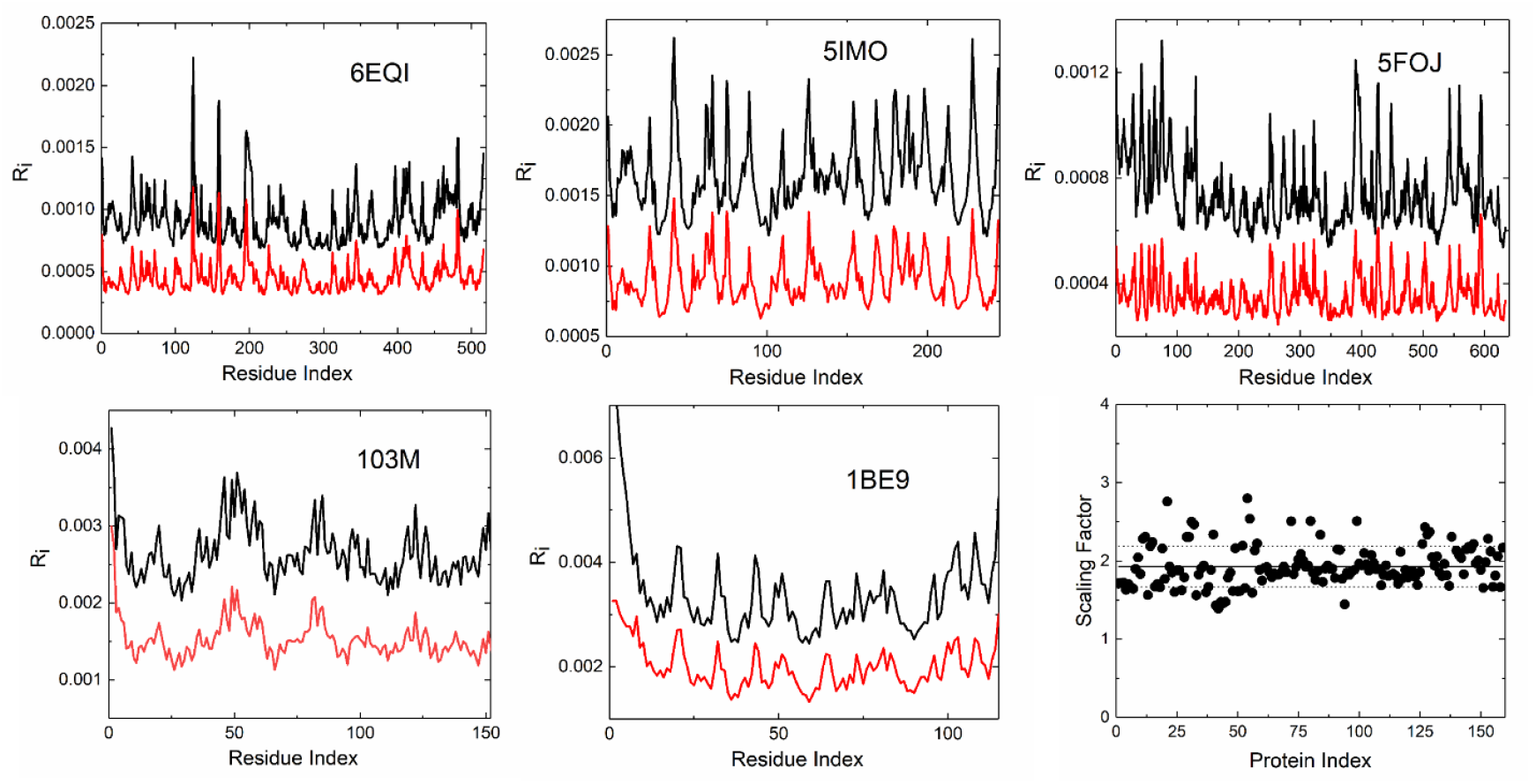
The top three and bottom two left panels show representative profiles of *R*_*i*_ (mean dynamic distance, black lines) and *R*_*i,edge*_ (mean dynamic distance to direct neighbors, red lines) versus residue index for five proteins from the dataset. The parallel curves with *R*_*i*_ = *CR*_*i,edge*_ illustrate that long-range interactions contribute equally to local neighbor interactions in determining total dynamic distance. The bottom right panel shows the distribution of the scaling coefficient *C* in the relationship *R*_*i*_ = *CR*_*i,edge*_ across 200 non-helical proteins. The horizontal solid line indicates the mean value (C = 1.93 ± 0.02), and dotted lines mark ±1 standard deviation. The near-zero slope confirms a universal scaling relationship for mixed and β-sheet structures, while all-helical proteins deviate from this pattern and are excluded from the analysis. All-α-helical proteins (excluded from fit) systematically deviate beyond ±3 SD (not shown), demonstrating structure-dependent scaling regimes. The Python code for calculating *R*_*ij*_ and *R*_*i*_ is given in the repository https://github.com/burakerman/spanning_tree_1.

Screening more than 200 diverse proteins revealed that all structures beyond ±3 SD from the mean were exclusively all-α-helical proteins. This systematic deviation reflects the anisotropic geometry of helical bundles, where extended secondary structures create directionally constrained communication pathways. The anisotropic, quasi-1D connectivity of helices may decouple local neighbor motions from the global network response, violating the isotropic coupling assumption implicit in the scaling. All 200 mixed α/β and β-sheet proteins showed C = 1.93 ± 0.02, demonstrating universal local-to-global coupling for compact, isotropic architectures.

The linear correspondence between *R*_*i*_ and *R*_*i,edge*_ holds within each protein structure: for a given protein, *R*_*i*_ = *CR*_*i,edge*_ across all residues, as evidenced by the parallel curves in the representative profiles. The scaling factor *C* is remarkably consistent across diverse protein architectures, with a mean value of 1.93 ± 0.02 over 200 non-helical proteins. This near-universal proportionality implies that functional roles are preserved across scales within a given protein. Residues identified as bottlenecks (large *R*_*i,edge*_) retain their bottleneck character globally (large *R*_*i*_), and locally redundant edges (small *R*_*i,edge*_) remain redundant globally (small *R*_*i*_). Hence, these residue classifications are effectively scale-invariant, maintaining their structural–functional identity independent of whether local or global measures are used.

The consistency of the scaling factor shows a fundamental local-to-global coupling principle: within each protein, a residue’s local dynamic properties directly determine its global properties through an approximately fixed proportionality. This arises because both *R*_*i*_ and *R*_*i,edge*_ are governed by the same underlying dynamical architecture encoded in the Laplacian pseudoinverse. The factor of approximately 2 indicates that non-local interactions contribute roughly equally to local interactions in determining total dynamic distance. This local-to-global coupling demonstrates that a residue’s immediate neighborhood properties propagate throughout the entire protein through the network’s dynamical structure, linking local dynamics and global allostery through a simple, nearly universal linear relationship.

In the following, we use *R*_*ij*_ and *R*_*i*_ as the fundamental observable that link network topology to information dynamics. By analyzing communication pathways across proteins and within specific systems such as KRAS, we demonstrate that multiplicity of parallel pathways enables intrinsic error correction within the protein’s dynamic architecture.

### B. Edge and path probabilities

Motivated by the ensemble nature of intramolecular communication introduced above, we treat allosteric signal propagation as a statistical process over multiple possible paths. Quantifying this process requires assigning probabilities to individual communication paths.

When a residue is displaced, by ligand binding, mutation, or thermal fluctuation, this perturbation spreads across the network on short time scales. Any mathematical description of information transfer must therefore account for global connectivity. A spanning tree captures a minimal set of couplings sufficient to transmit a perturbation throughout the protein. Formally, a spanning tree is a subnetwork that includes all residues (nodes), contains no cycles, and uses the minimal number of edges required to maintain connectivity. While perturbations may propagate through many possible routes in the full contact network, each such route can be associated with an underlying spanning tree that specifies its essential connectivity. Spanning trees thus provide a natural mathematical representation of the fundamental skeletons through which dynamic information propagates.

Here, we consider the uniform distribution over all spanning trees [36-38]: every spanning tree of the network is assigned equal probability. Sampling from this distribution produces an ensemble of minimal, cycle-free network backbones. Different spanning trees emphasize different alternative communication routes, and the statistics of this ensemble characterize how strongly a given residue pair depends on particular connections.

Once the uniform spanning–tree measure is defined, two natural probabilistic quantities become biologically informative:

1. Edge probability. For any edge (*i, j*), one may compute the probability, *P* (*i, j*), i.e., the fraction of spanning trees that contain (*i, j*) or equivalently the probability that (*i, j*) is an edge of a randomly chosen uniform spanning tree. Edges with high spanning-tree probability are structurally important hubs, used by many spanning trees, and lie in bottleneck regions with few alternative routes, making them highly perturbable; in contrast, edges with low probability are redundant, used by few spanning trees, located in rigid, well-connected regions with many alternative pathways, and thus least affected by perturbations. This confers a structural redundancy that allows them to maintain communication and remain largely unaffected by perturbations. In Appendix B we show that

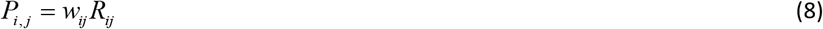

Thus, the dynamic distance landscape highlights residues that are likely to play key roles in allosteric communication.
2. When residues i and j are not directly linked by an edge in the network, each spanning tree provides a unique path connecting residues i and j, so that the total number of possible communication routes equals the total number of spanning trees of the network. In practice, however, the contribution of these paths is not uniform: a small subset of topologically or dynamically favorable routes carries the majority of the probability weight. Thus, while the spanning-tree ensemble provides a complete basis for all possible communication channels, only a few dominant paths are effectively used in the dynamics. High-probability paths are those that lie in topological bottlenecks of the structure, or in dynamically important regions where alternatives are scarce. The probability *P* = *P*(*v*_1_, *v*_2_,…, *v*_*m*_) that a given spanning tree contains the given path *v*_1_, *v*_1_,…, *v*_*m*_, which is the frequency of occurrence of that path in all spanning trees is given in Appendix D. The Transfer-Current theorem due to Burton and Pemantle provides the statistical weight of specific edge configurations between two nodes in random spanning tree ensembles [39,40]. For the simple path *P* = *P*(*v*_1_, *v*_2_,…, *v*_*m*_) connecting source *s* = *v*_1_ to target *t* = *v*_*m*_, with edge set *E*_*P*_= {*e*_1_, *e*_2_,…, *e*_*m*−1_}, we define the edge response matrix *K*_*E*_ which is a function of the elements of *K* . *K*_*E*_ is of size (m−1)×(m−1) with elements:

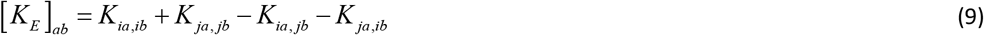

where *e*_*a*_ = (*v*_*ia*_, *v* _*ja*_) and *e*_*b*_ = (*v*_*ib*_, *v* _*jb*_) (See Appendix D for the detailed derivation of the elements of [*K*_*E*_ ]_*ab*_). The absolute probability that all edges in P are simultaneously present as the unique connecting path between s and t in a random spanning tree is:

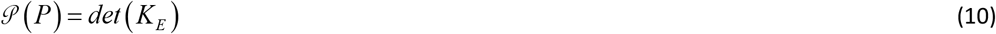

This result arises from considering all spanning trees of the graph and measuring the probability that all edges in path P belong to a uniformly random spanning tree [29]. Given the set of m simple paths *P*_*s*→*t*_ = {*P*_1_, *P*_2_,…, *P*_*m*_} connecting residues s and t, we compute relative weights:

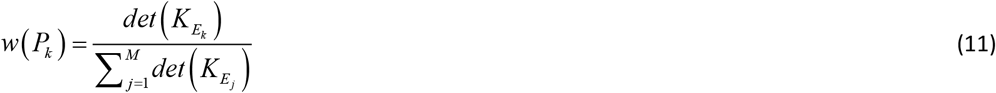

where the denominator sums over all m paths between the specified endpoints. These weights *w* ( *P*_*k*_) represent the likelihood that, among all possible connections between s and t, the communication follows path *P*_*k*_ .

Other quantities, such as the distribution of path lengths and the expected number of shared edges between distinct residue pairs, can be computed using the same framework. These provide additional structural information about redundancy and bottlenecking in the protein’s communication architecture.

### C. Information transfer, channel multiplicity and noise filtering

The dynamic distance *R*_*ij*_ not only quantifies residue fluctuations but also captures how information propagates through the protein network. Because *R*_*ij*_ reflects, due to the local to global scaling property, the ensemble of pathways connecting residues i and j, it inherently captures the multiplicity of spanning trees between these residues. This multiplicity underlies two complementary forms of intrinsic error correction. First, random fluctuations along different paths tend to cancel, reducing noise in the transmitted signal. Second, if a perturbation is partially blocked or attenuated along one route, alternative pathways continue to transmit the signal, maintaining the fidelity of information transfer. Thus, the dynamic distance acts as an integrated measure of both the diversity and the reliability of communication routes within the protein.

To illustrate how multiplicity suppresses noise, consider a perturbation transmitted simultaneously through N independent, equally probable pathways. Let the signal arriving along each path consist of the true signal plus a random fluctuation with variance *σ* ^2^ . In Appendix C, we show that when the outputs of these paths are combined at the target residue, the independent fluctuations average out, and the net variance of the received signal becomes

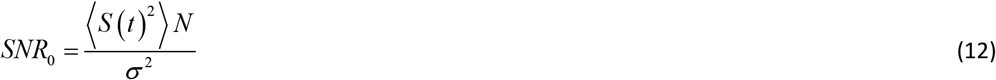

where *SNR*_0,*ij*_ is the signal to noise ratio for information transfer between nodes i and j, and ⟨*S* (*t*)^2^ ⟩ is the input signal power. For comparative purposes, the normalization ⟨ *S* (*t*)^2^ ⟩ / *σ* ^2^ = 1 may be used, which we adopt in calculations here. This normalization simply fixes the unit of signal-to-noise ratio and is equivalent to choosing a reference noise scale. All reported results depend only on relative SNR values and are invariant under rescaling.

Thus, the more distinct pathways contribute, the smaller the impact of random noise on the overall transmitted signal. In this sense, multiplicity acts as a natural noise filter: many weakly noisy routes together produce a more reliable readout than any single route alone [41].

When the paths are not of equal probability, use of the Burton-Pemantle theorem (Appendix E) and obtain

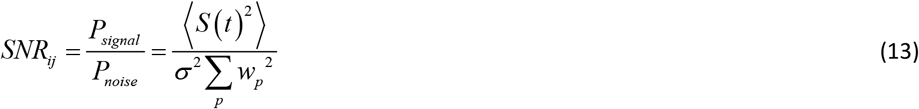

where *SNR*_*ij*_ is the signal to noise ratio for information transfer between nodes i and j, and *w*_*p*_^2^’s are obtained from Eq. 11. If the probabilities are uniform, i.e., *w*_*p*_ = 1/ *N* then Eq. 13 reduces to Eq. 12. By systematically analyzing the distribution of *R*_*ij*_ across a protein, one can identify regions that act as critical hubs for information processing and regions that act as flexible buffers, together implementing intrinsic error-correcting mechanisms.

The distinction between uniform and network-derived pathway probabilities is essential for interpreting signal-to-noise ratios in protein communication. In the uniform case, all communication pathways contribute equally, corresponding to a maximally delocalized ensemble in which no single route is privileged. This represents an idealized upper bound on pathway multiplicity and therefore on noise suppression. In contrast, Burton–Pemantle probabilities account for the underlying network topology and elastic couplings, assigning higher weight to pathways that occur more frequently across the spanning-tree ensemble. As a result, communication becomes heterogeneous, with signal transmission concentrated along a subset of dominant routes. This concentration reduces effective pathway multiplicity and lowers the signal-to-noise ratio relative to the uniform case.

We note that *SNR*_0_ (uniform) is an upper bound representing maximal possible noise suppression, while *SNR* (topology-weighted) is the physically realized value. The difference *SNR*_0_ − *SNR* quantifies the “cost” of funneling signal through preferred paths.

This distinction can be quantified using Rényi entropy, which measures the effective number of contributing pathways. For a set of pathway probabilities { *p*_*α*_ }, the order-2 Rényi entropy,

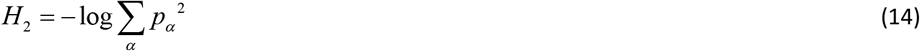

directly reflects the degree of pathway concentration: it is maximal for uniform probabilities and decreases as communication becomes dominated by fewer high-probability paths. Consistent with the SNR expressions derived above, lower Rényi entropy corresponds to reduced noise averaging and hence lower SNR. Thus, comparing uniform and Burton–Pemantle probabilities provides a principled way to distinguish idealized, maximally redundant communication from the topology-constrained information transfer realized by the protein. In the Results section, we demonstrate how this entropy reduction varies across proteins and relates to their allosteric architectures.

### D. Entropy perturbability as a physical basis for dynamic distance

Although the entropy-perturbability relations derived below are not used explicitly in the path multiplicity calculations, they provide the physical basis for the dynamic distance metric and for interpreting bottleneck and redundant interactions. In particular, they clarify how local changes in network connectivity can induce global responses in protein dynamics.

Coupling between local and global degrees of freedom can be analyzed in terms of entropy perturbability. In Appendix A, we derive the relation

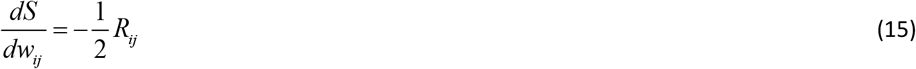

where an increase in the weight, *w*_*ij*_, of the edge ij results in a decrease in entropy proportional to the dynamic distance, *R*_*ij*_ . Thus, a change in the strength of a bottleneck edge causes a larger change in entropy than a change in a redundant edge.

If all edges incident to a residue p are perturbed simultaneously, the resulting entropy change follows an analogous expression involving the summed dynamic distances to that residue (Appendix A). This establishes a direct connection between residue-level perturbations and global entropic response.

This framework introduces an effective, perturbation-induced directionality in the response of the system. Residues whose perturbations generate large system-wide entropy changes act as entropy sources, driving fluctuations through correlated motions, whereas residues whose perturbations lead to minimal global entropy shifts act as entropy sinks, effectively absorbing fluctuations. Bottleneck residues, identified by their high sensitivity to unit perturbations, behave as entropy sources due to their role in channeling communication through a limited set of high-probability pathways. Conversely, redundant residues behave as entropy sinks because pathway multiplicity buffers their local fluctuations, yielding small net entropy changes.

Although the underlying network couplings are reciprocal, this source–sink asymmetry emerges from how perturbations are distributed across the ensemble of communication pathways, as quantified by dynamic distance and path probabilities. This perspective is conceptually related to transfer entropy approaches, which infer directionality from system responses to perturbations rather than from static connectivity alone [42]. These response asymmetries are illustrated explicitly in the Results section using KRAS as a representative case study.

## III. Results

As a comprehensive demonstration of the principles of dynamic information processing, including path probabilities, pathway multiplicity, intrinsic error correction, and entropy-driven signal propagation, we first apply the Gaussian Network Model to the oncogenic protein KRAS. This detailed analysis provides topology-driven insights into KRAS allostery and serves as a concrete illustration of the theoretical framework developed above.

We then extend the analysis to a benchmark set of 20 proteins with experimentally characterized allosteric and orthosteric sites, focusing on the distribution of communication channel probabilities and their deviation from uniform-path models.

### A. Bottleneck and redundant residues of KRAS

We first computed the full matrix of dynamic distances *R*_*ij*_ using Eq 3 for all residue pairs. From these, using Eqs. 5 and 6, we calculated *R*_*i*_ and *R*_*i,edge*_ shown in Figure 3.

**Figure 3.**
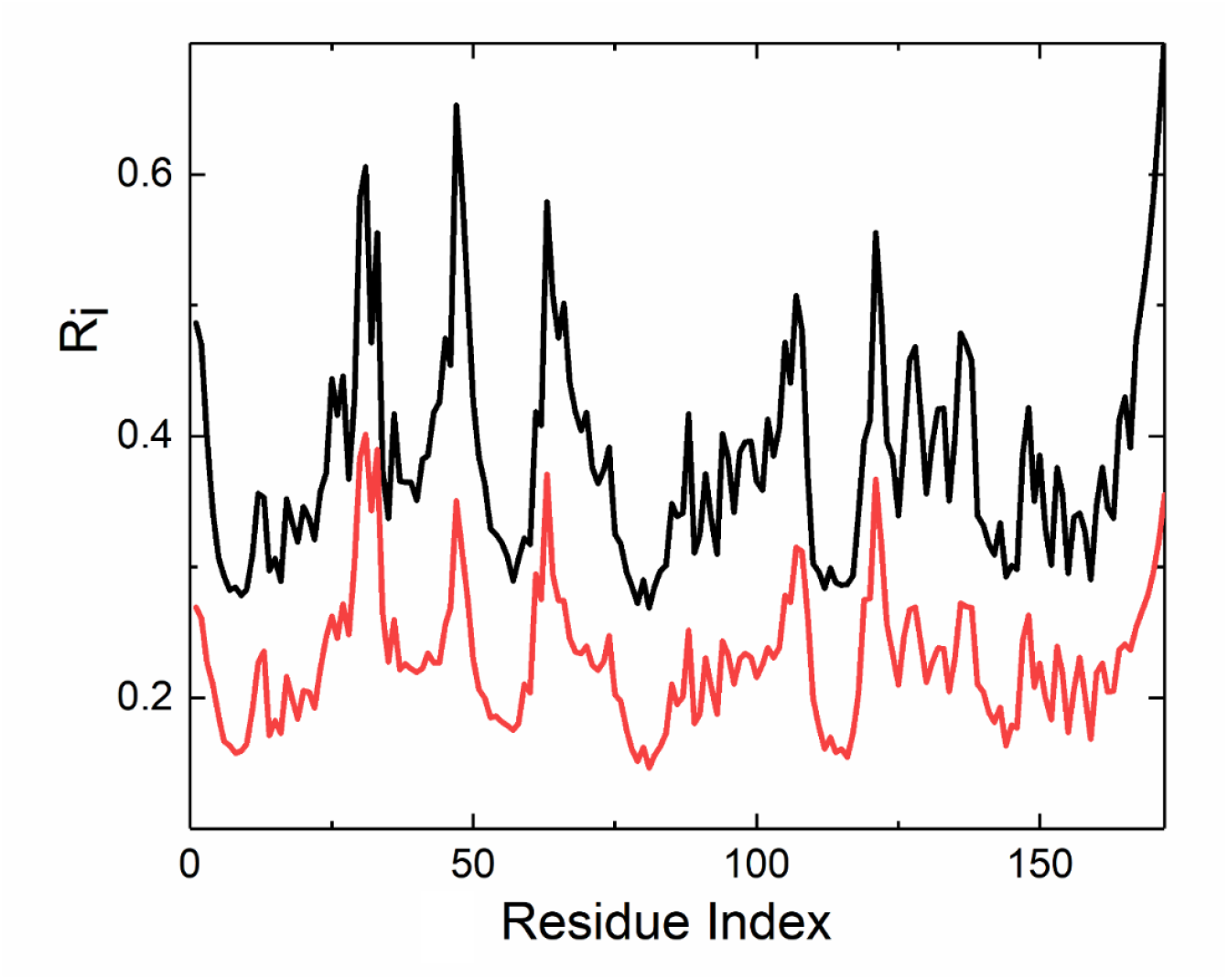
Residue dynamic distance profiles obtained from the Gaussian Network Model (GNM) with a cutoff distance of 7.8 Å for 6GOD.pdb. The black and red lines represent the global residue response *R*_*i*_ and the edge-averaged response *R*_*i,edge*_, respectively, plotted as a function of residue index. Peaks correspond to bottleneck residues, while minima indicate redundant residues. The near-parallel behavior of the two curves supports the validity of Eq. 7 and allows estimation of the scaling parameter C as 1.75.

The peaks and dips of the curves align with residues that have been experimentally identified as structural allosteric hubs involved in nucleotide binding and effector interactions, consistent with their proposed role in maintaining robust functional information flow [43-49]. A cutoff distance of 7.8 Å corresponds to the first coordination shell of C^α^ atoms, capturing direct residue-residue contacts. This value was optimized in previous GNM studies [50] to balance connectivity and computational efficiency.

### B. Path probabilities and error correction in KRAS: The Q61-V114 path

Q61, located in Switch II (residues 60-76, β3 strand), serves as a catalytic bottleneck positioning the nucleophilic water for GTP hydrolysis, with GAP-stimulated arginine finger stabilizing Q61-G60 interactions. RAF binding near Q61 quenches Switch II fluctuations via H-bond networks (Q61-T35-G60-γ-phosphate), rigidifying the active conformation. Mutational atlases confirm Q61’s high betweenness centrality in β-sheet propagation, with perturbations yielding strong directional effects on RAF binding [45,47,51-53].

V114 (β5 strand, residues 110-116, allosteric lobe α3-α5 interface) functions as a redundant hydrophobic packer in network hubs linking cryptic pockets (H95/Y96) to effector sites, exhibiting low betweenness with multiple bypass paths via β4 parallels (F78/L80). MD and GNM analyses position V114 in resilient β-sheet transmission, where the protein network can adjust to mutations at V114 by using alternate pathways or interactions to maintain function. RAF binding propagates ∼30Å through the beta sheet, β3→β4→β5 path, tightening V114 to lock KRAS in closed conformation by allosteric coupling.

High-probability paths between Q61 (bottleneck) and V114 (redundant) traverse the central β-sheet, validated by more than 22,000 mutational scans showing strand-by-strand decay (strongest β3, diminishing to β5). RAF-induced Q61 quenching anticorrelatedly reduces V114 entropy, stabilizing hydrophobic packing to prevent premature dissociation. Path calculations show dominant β-sheet routes matching this experimental coupling.

Our analysis shows that allosteric communication between Q61 and V114 occurs not through a single predetermined route, but through a statistical ensemble of pathways. Using the Burton-Pemantle theorem applied to protein contact networks (|See Eq. 11 and Appendix D), we computed the probability distribution over all simple paths connecting the two allosteric sites. We identified 5461 distinct pathways of lengths 4-7 residues, each assigned a statistical weight:

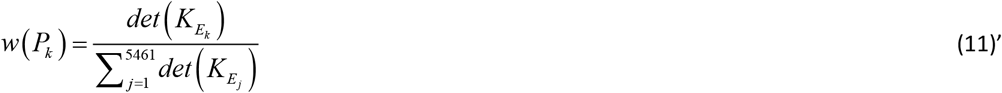

These normalized weights satisfy 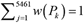 and represent the relative likelihood that each pathway serves as the primary conduit for signal transmission. The maximum probability paths obtained according to Eq. 11’ are shown in Figure 4.

**Figure 4.**
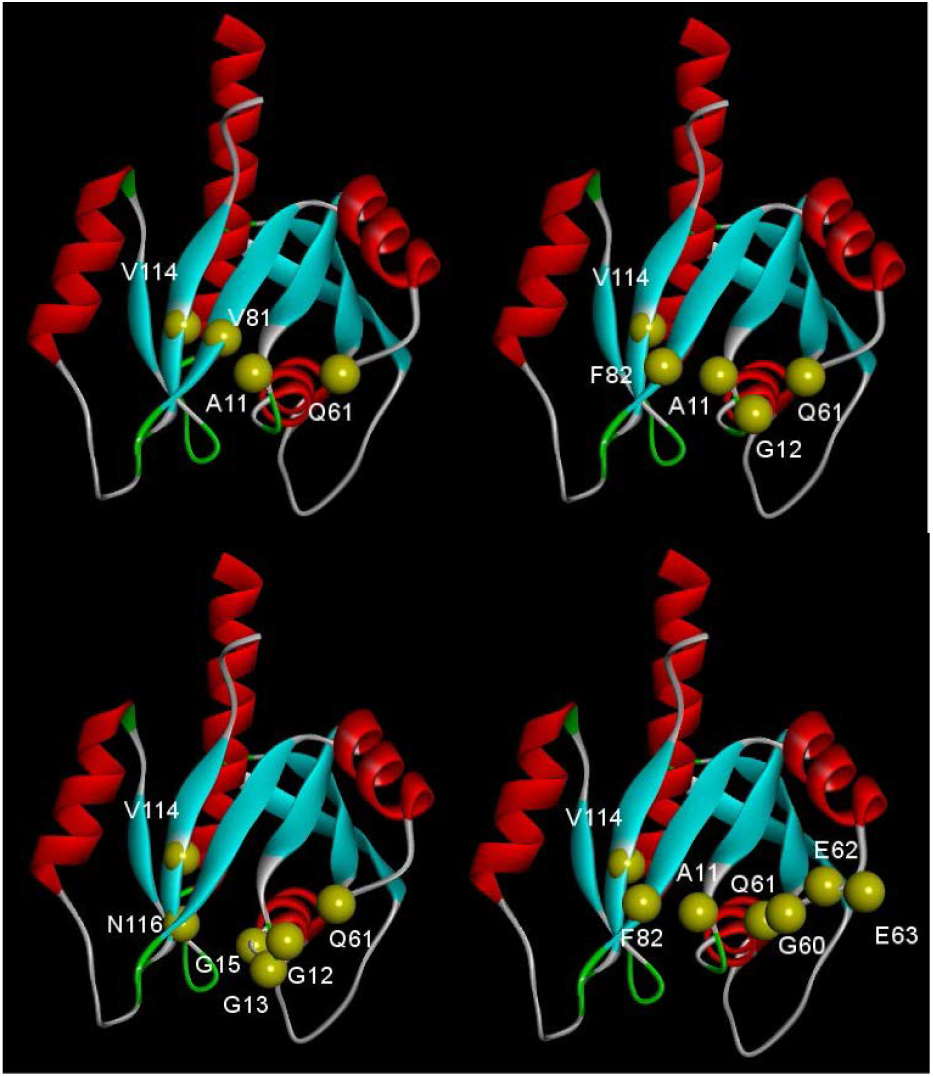
Highest-probability communication paths between Q61 and V114 in KRAS, PDB file: 6GOD.PDB, Cutoff distance: 7.8 Å. Four representative paths of increasing residue path length are shown: (top left) length 4, (top right) length 5, (bottom left) length 6, and (bottom right) length 7. For clarity, Helix 1 has been removed to improve the visibility of the connecting routes. Residues along each path are displayed as yellow spheres and labeled with their identities and sequence numbers. The specific paths are: Path 1 (length 4): Q61 → A11 → V81 → V114 Path 2 (length 5): Q61 → G12 → A11 → F82 → V114 Path 3 (length 6): Q61 → G12 → G13 → G15 → N116 → V114 Path 4 (length 7): Q61 → E63 → E62 → G60 → A11 → F82 → V114.

The signal to ratio values are: SNR_0ij_= 5461.0, SNR_ij_= 160.8. The ratio of SNR_ij_ to SNR_0ij_ is the efficiency and is 2.9%. This low efficiency indicates that KRAS’s topology strongly concentrates signal flow, sacrificing potential noise suppression for specific, high-fidelity routing through its β-sheet core. The order-2 Rényi entropy, H2, for uniform probability paths is 12.415 bits, and the actual H2 is 10.384 bits with an entropy gap of 2.031 bits.

This distribution shows two crucial insights: First, while longer paths are numerically more abundant, individual shorter paths have dramatically higher probabilities. From column 3 of Table 1 we see that a typical 4-residue path is approximately 650 times more likely to transmit signal than a typical 7-residue path, an exponential decrease with path length. Second, the system employs a multi-scale signaling strategy: a few high-probability direct routes for efficient communication, complemented by thousands of low-probability alternative pathways that provide redundancy and robustness.

**Table 1.**
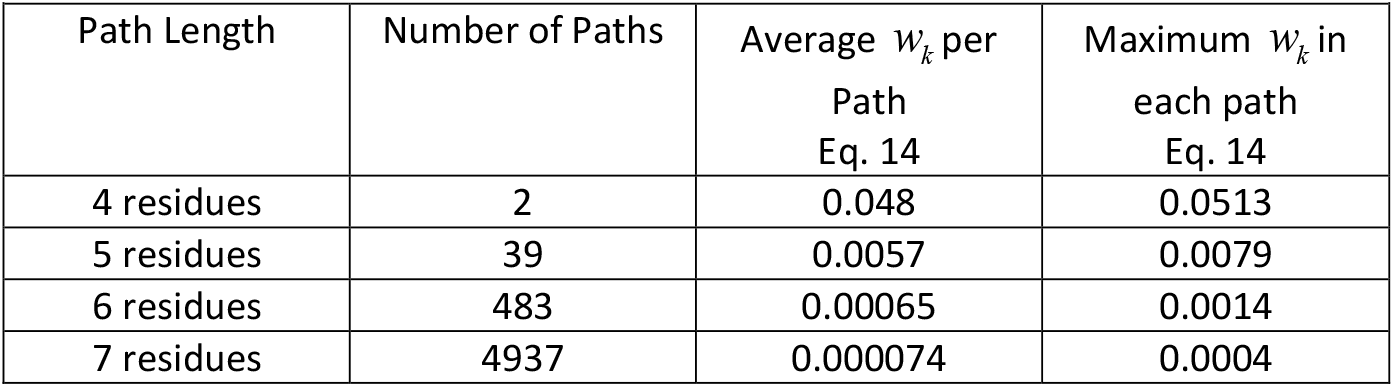
Dominant Pathways and Pathway Length Distribution.

Paths were limited to a maximum length of seven residues based on physical constraints, statistical insignificance of longer paths (<0.1% probability weight), and consistency with experimentally characterized allosteric routes. This cutoff captures more than 99.9% of statistically relevant pathways while remaining computationally tractable.

Despite the large number of longer pathways (e.g., thousands of paths of length 7), they play a negligible role in determining the signal-to-noise ratio (SNR). This follows directly from the fact that the SNR is inversely proportional to the sum of squared path weights, 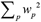. Although the number of available paths grows rapidly with path length, the probability assigned to each individual path decays even more strongly. In the present data, each additional residue in a path reduces the average path weight by approximately an order of magnitude; when squared, this results in a strong suppression of the contribution of long paths to 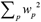. Consequently, the SNR is dominated by a small number of short, high-probability pathways, while longer pathways primarily increase pathway multiplicity without significantly affecting signal fidelity.

### C. Path probabilities and error correction in various allosteric proteins

We studied 20 allosteric proteins belonging to different functional classes; the results are summarized in Table 2. The signal-to-noise ratio spans a wide range across these systems. While some proteins show comparable values of SNR_0_ and SNR, others exhibit a substantial enhancement upon Burton–Pemantle weighting. The Rényi-2 entropy reduction H_20_-H_2_ also varies widely, with larger values indicating that signal transmission is dominated by a small number of short, high-probability pathways, whereas longer paths primarily contribute to noise suppression.

**Table 2.** SNR values for 20 allosteric proteins.

| Table 2. SNR values for 20 allosteric proteins |  |  |  |  |  |  |  |  |  |
| --- | --- | --- | --- | --- | --- | --- | --- | --- | --- |
| Protein | Orthosteric | Allosteric | PDB | Total No. paths=<br>SNR<sub>0</sub> | SNR | ΔSNR% | H<sub>20</sub> (bits) | H<sub>2</sub> (bits) | Entropy gap (bits) |
| DHFR [54] | D27 | G121 | 1RX2 | 1705 | 215.2 | 12.6 | 10.736 | 9.526 | 1.209 |
| PDZ3 [55] | G330 | I327 | 1BE9 | 17828 | 502 | 2.8 | 14.122 | 11.867 | 2.255 |
| Src kinase [56] | K295 | E310 | 2SRC | 1611 | 331.2 | 20.6 | 10.654 | 9.869 | 0.785 |
| β2AR [57] | D113 | R131 | 2RH1 | 492 | 232 | 47.2 | 8.943 | 8.573 | 0.370 |
| Hemoglobin [58] | H87 | H146 | 2HHB | 3354 | 120.7 | 3.6 | 11.712 | 9.227 | 2.484 |
| Adenylate K [59] | D93 | R156 | 4AKE | 25 | 23.6 | 94.4 | 4.644 | 4.595 | 0.049 |
| IGPS [60] | C84 | H178 | 1GPW | 109 | 60.2 | 55.2 | 6.768 | 6.433 | 0.325 |
| Hsp90 [61] | E33 | R380 | 2CG9 | 265 | 42.4 | 16.0 | 8.050 | 7.119 | 0.931 |
| LacI [62] | H74 | D274 | 1LBG | 237 | 41.5 | 17.5 | 7.889 | 6.953 | 0.935 |
| ATCase [63] | D236 | E239 | 1AT1 | 529 | 20.9 | 4.0 | 9.047 | 5.861 | 3.186 |
| p53 [64] | R248 | R175 | 1TSR | 13834 | 144.2 | 1.0 | 13.756 | 10.746 | 3.010 |
| CAP [65] | E72 | S62 | 1G6N | 1131 | 169 | 15 | 10.143 | 9.195 | 0.949 |
| CheY [66] | D57 | Y106 | 3CHY | 12026 | 61.6 | 0.5 | 13.554 | 9.507 | 4.047 |
| Adenosine A2A [67] | N253 | Y271 | 5UIG | 3018 | 187.4 | 6.2 | 11.559 | 9.877 | 1.683 |
| Glutathione S-T [68] | Y7 | H50 | 3D0Z | 6546 | 252.7 | 3.9 | 12.676 | 10.555 | 2.121 |
| PDK1 Kinase [69] | D146 | L155 | 4A07 | 6787 | 48.2 | 0.7 | 12.729 | 9.251 | 3.477 |
| Caspase-1 [70] | C285 | R286 | 1RWK | 4777 | 37.1 | 0.8 | 12.222 | 7.545 | 4.677 |
| Muscarinic M2 [71] | D103 | W422 | 4MQT | 389 | 73.1 | 18..8 | 8.604 | 7.736 | 0.867 |
| Adrenegic β2 [67] | D113 | Y316 | 5X7D | 2384 | 58.4 | 2.5 | 11.219 | 8.699 | 2.520 |
| mGluR5 [72] | S809 | Y659 | 4O09 | 3513 | 87.2 | 2.5 | 11.778 | 9.896 | 1.882 |
| SNR<sub>0</sub>: signal to noise ratio for uniform probabilities, SNR: signal to noise ratio calculated from Burton Pemantle theory, ΔSNR: difference between uniform and Burton-Pemantle probabilities, H<sub>20</sub>: Renyi entropy of order 2 calculated using uniform probabilities, H<sub>2</sub>: calculated using Eq. 13, Entropy gap=H<sub>20</sub>-H<sub>2</sub> | | | | | | | | | |

We observed no systematic trend linking communication efficiency metrics (e.g., entropy gap, ΔSNR%) to broad functional categories (enzymes, kinases, GPCRs). For instance, enzymes in our set exhibited both high (ATCase) and low (Adenylate Kinase) ΔSNR values. This variability indicates that the statistical architecture of allosteric communication is a property of the individual protein’s network topology, rather than a feature conserved across functional classes.

## Discussion

The idea that multiple allosteric routes enhance robustness has been discussed qualitatively [6], but a quantitative link between pathway multiplicity and noise suppression has been lacking. Here we provide such a framework by combining spanning-tree statistics with a network description of residue connectivity.

Our analysis of KRAS serves as a proof of concept, showing that spanning-tree statistics yield exact pathway probabilities, quantify intrinsic noise filtering, and distinguish critical from redundant contacts. The approach is readily generalizable to other proteins.

Rather than discovering allosteric pathways *de novo*, our method quantifies the statistical ensemble of communication routes between pre-specified functional residues. Given a source residue sss and target residue ttt, we compute the exact probability distribution over all simple connecting paths using the Burton–Pemantle theorem. This conditional analysis identifies dominant routes, quantifies pathway multiplicity and noise filtering, and distinguishes critical from dispensable contacts via resistance (dynamic) distances. While functional sites must be specified externally, this reflects biological reality, where allostery occurs between defined functional locations.

Unlike heuristic centrality measures, the spanning-tree framework assigns exact statistical weights to pathways and quantifies edge redundancy through resistance distances, providing a physically grounded alternative to phenomenological network metrics. Both pathway importance and redundancy emerge naturally from the graph Laplacian.

Dynamic distance provides a mechanical link between network topology and allosteric communication. Small values indicate strong coupling and high redundancy, while large values identify bottleneck edges whose perturbation strongly affects network entropy.

For KRAS, the analysis predicts large pathway multiplicity and substantial noise reduction arising from parallel connectivity alone. If general, such topological features may explain how proteins maintain robust allostery despite sequence variation and thermal noise, indicating that network architecture contributes intrinsic signal-processing capability. The systematic exclusion of helical proteins from the universal scaling regime shows that local-to-global coupling depends on overall protein architecture. In all-α proteins, the extended geometry of helices creates anisotropic contact networks where longitudinal (along helix) and transverse (between helices) couplings differ substantially. In contrast, β-sheets and mixed architectures exhibit more isotropic contact distributions, yielding universal scaling independent of protein size, function, or evolutionary origin.

More generally, network topology organizes dynamic fluctuations into multiple communication routes that act as error-correcting mechanisms. Residues connected by many low-resistance paths form robust communication hubs, preserving global function against thermal or mutational perturbations.

Several limitations remain. The model treats all contacts equally. This ignores chemical specificity, hydrophobic cores, salt bridges, and H-bonds, which will drastically alter true “channel” quality.This is a topological prior, and future work should integrate energetic weights ( *w*_*ij*_ ≠ 1). Functional validation therefore requires integration with mutagenesis, functional assays, evolutionary analysis, and dynamic measurements. Mutations correspond to changes in contact topology and Laplacian structure and are expected to reorganize dynamic-distance profiles, shifting bottleneck and redundant regions. Quantifying these effects is a natural direction for future work.

## Conclusion

Proteins are dynamic systems in which internal fluctuations transmit functional signals through structured networks of residue contacts. Using spanning-tree statistics, we show that network topology alone organizes these fluctuations into multiple communication pathways that provide intrinsic noise suppression.

For KRAS, thousands of parallel paths contribute to allosteric communication, yielding a ∼200-fold noise reduction while concentrating signal transmission onto a small number of statistically dominant short routes. This architecture explains how robust allosteric signaling can be maintained despite thermal noise and sequence variability.

More generally, our framework provides a quantitative link between protein structure, dynamics, and information flow, demonstrating that error-correction–like behavior emerges naturally from network connectivity. These results suggest that robustness in protein allostery arises from topological organization rather than fine-tuned interactions, and they offer a foundation for extending information-theoretic concepts to protein dynamics and biomolecular design.

Repository address: https://github.com/burakerman/spanning_tree_1

## Appendix A Entropy perturbability. Local to global mapping

*L* is the Laplacian defined by Eq. 2 and *K* = *L*^+^ is the Moore-Penrose inverse of the Laplacian, we can write:

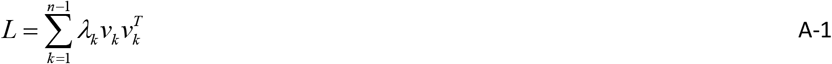

where *λ*_*k*_ > 0 are the non-zero eigenvalues (we exclude the zero eigenvalue corresponding to the constant eigenvector) and *v*_*k*_ are the corresponding eigenvectors.

Then:

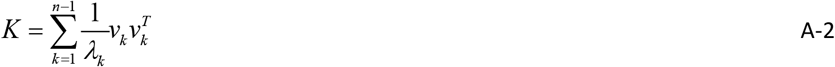

The (differential) entropy for a Gaussian random vector with covariance matrix *K* is given by

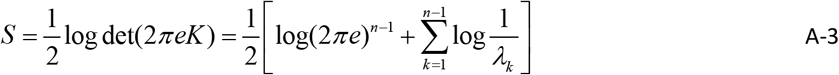

Differentiating *S* with respect to the weight *w*_*pq*_ of an edge

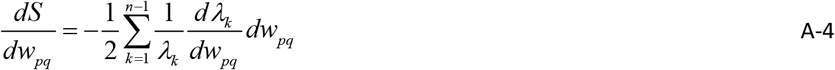

We then perturb edge weight *w*_*pq*_ by δ:

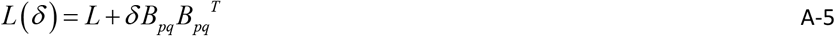

where *B*_*pq*_ *=e*_*p*_ − *e*_*q*_ with the basis vector *e*_*p*_ being a vector with 1 at location p and zero elsewhere. By first-order perturbation theory:

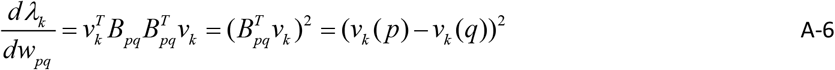

Substituting back in Eq. A-4

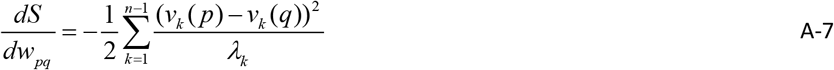

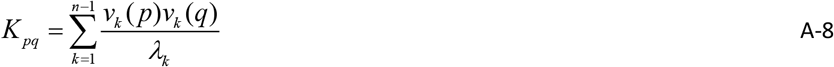

and:

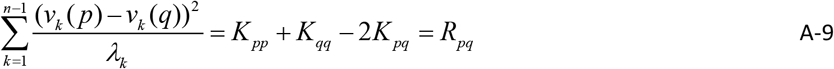

Eq.A-7 then becomes

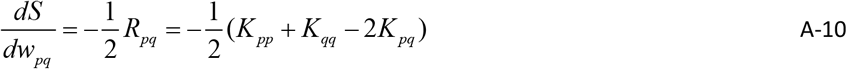

This relation shows that a local perturbation of an individual edge weight is directly coupled to a global property of the network, the entropy. In particular, changes in an edge weight, such as those induced by a mutation, produce an entropy variation proportional to *R*_*ij*_ . As shown in Appendix B, *R*_*ij*_ admits a probabilistic interpretation as the probability that residues i and j are connected by an edge in a spanning tree of the network.

Suppose all edges pendant to a node p are perturbed by *ε* _*p*_ . Summing over the edges that are connected to node p gives

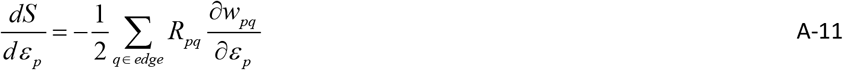

## Appendix B Derivation of dynamic distance from matrix-tree theorem and its relation to edge probabilities

Kirchhoff’s Matrix-Tree Theorem: Let *L*^*r*^ be the reduced Laplacian *L* (or “grounded” Laplacian) obtained by deleting the r-th row and r-th column from *L* . Its determinant, det *L*^*r*^, is equal to the number of all spanning trees of the graph, *τ* ( *L*) .

We represent a spanning tree by *T* and all spanning trees of the graph by 𝒯 In the context of the Matrix–Tree Theorem, *τ* ({*w*}) denotes the total weight of all spanning trees of the graph, defined as

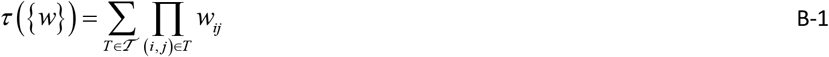

Expand *τ* ({*w*}) as a polynomial in the weights. The portion of*τ* proportional to *w*_*ij*_ equals the total weight of all spanning trees that include edge (i,j); denote that by *T*_(*i, j*)_ . Differentiating the polynomial extracts precisely those monomials:

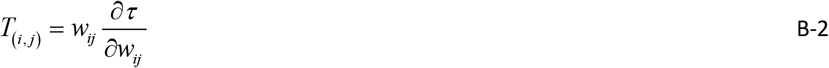

(Differentiation removes the factor *w*_*ij*_ from each monomial containing it; multiplying back by *w*_*ij*_ restores the full weight of those trees.)

The fraction of total spanning-tree weight that includes (i,j) is

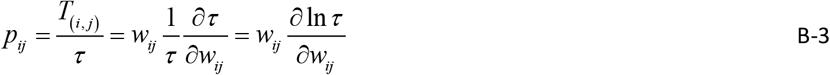

Changing a single edge weight *w*_*ij*_ modifies the Laplacian by the rank-one matrix

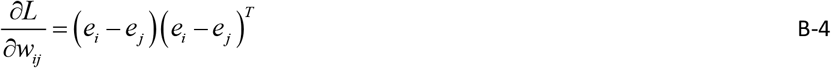

because off-diagonals *L*_*ij*_ = *L*_*ji*_ = −*w*_*ij*_ change and diagonals *L*_*ii*_ change by +*w*_*ij*_ . *e*_*i*_ is a vector with a 1 in the i-th position and 0’s elsewhere. Using *∂* ln det *M* / *∂M* = *M* ^−1^ for the cofactor minor and the fact ^1^ that the inverse of any cofactor is represented by the pseudoinverse on the nonzero subspace, one obtains

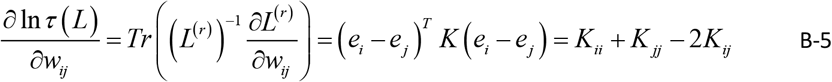

where, *Tr* ( *A*) id the trace of a matrix *A*, which is the sum of its diagonal elements. Combining B-3 and B-5 gives

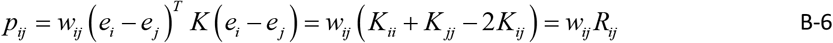

For the common unweighted case in GNM, *w*_*ij*_ = 1 for contacting residues, so

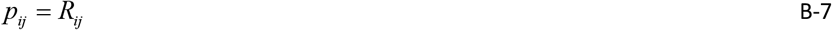

Thus, the dynamic distance between residues i and j equals the probability that their connecting edge appears in the spanning-tree ensemble. In this sense, *R*_*ij*_ quantifies how frequently communication between i and j is utilized across all possible global pathways.

## Appendix C Path multiplicity, time averaging, and signal-to-noise ratio with uniform path probabilities

When a perturbation applied at residue i propagates to residue j through multiple distinct structural pathways, the protein effectively implements a simple encoder–decoder architecture. The local perturbation is replicated into N parallel communication channels, corresponding to alternative paths through the residue network. Each channel transmits the same underlying signal but is subject to independent thermal and dynamical fluctuations. At the target residue, the physical dynamics act as a decoder that aggregates and time-averages the incoming signals from all channels. This collective averaging suppresses uncorrelated noise while preserving the coherent signal. As shown in the derivation below, the resulting effective noise variance is reduced by a factor 1/N, leading to a signal-to-noise ratio that increases linearly with the number of channels. Pathway multiplicity thus provides a quantitative mechanism by which proteins enhance the robustness of information transfer.

### 1. Encoder: identical signal injection

Let a time-dependent signal *S* (*t*) (e.g. a sinusoid applied at residue i) be copied identically into N physically distinct structural channels. The signal arriving through channel i is

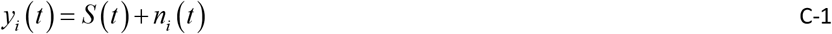

where, *S* (*t*) is the true input, *n*_*i*_ (*t*) is the noise added along channel i. In this section, all channels are assumed to be identical. We assume: *E* [ *n*_*i*_ (*t*)]= 0, *Var* [*n*_*i*_ (*t*)] = σ ^2^, *Cov* [*n*_*i*_, *n*_*i* ]_ = 0 *for i* ≠*∂j* (Independent zero-mean noise is the standard physical assumption for distinct structural paths.)

### 2. Decoder: time averaging of the incoming information

At the receiving residue j, the physical process is:

1. All incoming channel signals are summed:

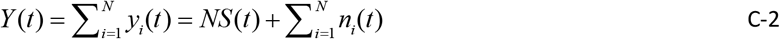
2. The system performs a time average because we are interested in the long-time, coarse-grained effect of the incoming fluctuations. In linear dynamical systems, this long-time response is mathematically equivalent to extracting the mean value of the sum of the incoming channel signals:

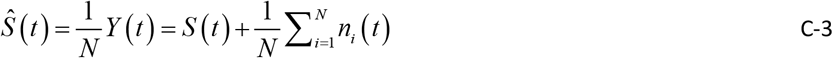

This is the decoder’s estimate of the signal. Thus the estimation error is:

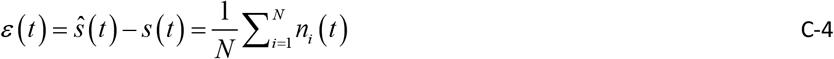
3. Noise reduction through path multiplicity

Because the noises *n*_*i*_ (*t*) are independent with variance _σ_ ^2^,

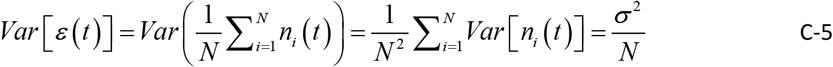

Therefore: noise power at the decoder scales as 1/N, noise amplitude (standard deviation) scales as 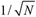, and signal-to-noise ratio *SNR*_0_ increases by a factor N. Here, the subscript 0 refers to uniform probabilities for the channels. Thus,

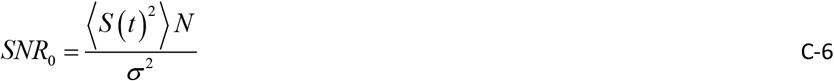

This establishes mathematically that an increase in the number of paths between i and j leads to stronger suppression of uncorrelated noise and, consequently, to more robust transmission of the signal to j. However, channels are not identical and independent because they are not equally utilized and there are several shared nodes and edges that lead to correlated noise. Thus, the proof above gives the upper bound, and real proteins achieve less due to the two factors above. Multiplicity of non-identical paths requires the development of path probabilities in Appendix D followed by the discussion of multiplicity in non-identical paths in Appendix E.

## Appendix D Path probabilities and the main code calculating these probabilities

This appendix describes the mathematical and computational methods implemented in the provided Python code for calculating path probabilities in protein contact networks using graph theory and random matrix theory. The method calculates the probability that all edges in a given path are simultaneously present in a uniformly random spanning forest of the protein graph.

### Input Data Processing

The method begins by parsing a Protein Data Bank (PDB) file using the Biopython PDB parser [73]. Only Cα atoms are considered, as they provide a simplified representation of the protein backbone. For each residue containing a Cα atom, the 3D coordinates are extracted and stored along with the residue identifier.

### Graph Definition

A protein contact graph G = (V, E) is constructed according to Eq. 2 where: vertices (V) represent Cα atoms of residues (nodes labeled by residue index), edges (E) connect two residues i and j if the Euclidean distance between their Cα atoms is ≤ *r*_*cutoff*_ (user-defined parameter, taken as 7.8 Å in this work in agreement with previous work [50])

This construction yields an undirected, unweighted graph representing the spatial proximity network of the protein structure.

### Graph Laplacian and Moore-Penrose Pseudoinverse

For a graph with n vertices, the combinatorial Laplacian matrix L is an n × n symmetric matrix defined by Eq. 2. Since the Laplacian of a connected graph is singular (rank n-1), we compute its Moore-Penrose pseudoinverse *K* = *L*^+^using the numpy.linalg.pinv() function, which employs singular value decomposition (SVD). The pseudoinverse satisfies: *LKL* = *L, KLK* = *K*, ( *LK*)^*T*^ = *LK*, ( *KL*)^*T*^ = *KL* .

### Burton-Pemantle or Transfer-Current Theorem for Path Probabilities [39]

The path probability calculation is based on the Transfer-Current Theorem from random matrix theory and electrical network theory [39,40]. For a path π consisting of edges e_1_, e_2_, …, e_k_, the probability that all these edges are simultaneously present in a uniformly random spanning tree is given by:

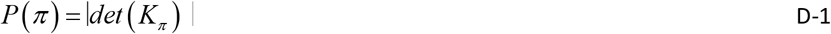

where *K*_*π*_ is a k × k matrix with entries:

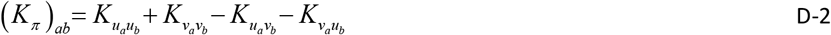

for edges e_a_ = (u_a_, v_a_) and e_b_ = (u_b_, v_b_), with K = L^+^ being the pseudoinverse of the graph Laplacian.

### Implementation

The code implements this theorem by:

1. Extracting all edges from a given path
2. Constructing the submatrix *K*_*π*_ according to the formula above
3. Computing the absolute value of the determinant of *K*_*π*_

This yields the raw probability for that specific path. We implemented a custom depth-first search algorithm to enumerate all simple paths of specific lengths between selected residues. The algorithm is implemented in the Python function ‘find_paths_of_length()’ (available in https://github.com/burakerman/spanning_tree_1). This function takes four inputs: the protein contact graph G, start residue s, end residue t, and desired path length 𝓁 (where 𝓁 = 3, 4, 5, 6, or 7 residues). The algorithm uses a stack-based iterative approach (rather than recursion) to avoid Python recursion limits. It maintains a stack of (current_node, current_path) tuples and explores all neighbors not already in the current path. Neighbors are sorted by residue index to ensure deterministic output ordering. The search terminates when paths reach the specified length 𝓁. We constrain path lengths to 3-7 residues based on biological considerations: allosteric communication typically involves 2-5 intermediate residues, and paths longer than 7 residues become combinatorially explosive while adding little biological relevance. This implementation choice differs from using NetworkX’s built-in ‘nx.all_simple_paths()’ function, allowing us to enforce exact length constraints and optimize memory usage for protein-sized graphs.

### Probability Normalization

Two normalization schemes are employed:

1. Within-length normalization: For paths of a specific length 𝓁, probabilities are normalized so that:

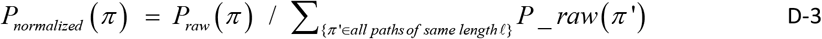

This makes probabilities sum to 1 for each path length separately.
2. Cross-length normalization: For comparison across different path lengths, all paths (lengths 3-7) are normalized together:

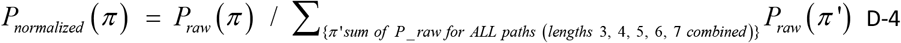

This makes the total probability across all paths of all lengths sum to 1.

### Output and Analysis

The code generates multiple output files:

paths_length_𝓁.txt: Contains all paths of specific length 𝓁 with within-length normalized probabilities path_length_summary.txt: Summary of relative contributions of different path lengths top_paths_analysis.txt: Analysis of the most probable individual paths

### Statistical Analysis

The code performs several analytical computations:

1. Relative contribution of each path length to total probability
2. Average probability per path for each length
3. Identification of top individual paths by probability
4. Comparison of top path probability versus average for each length

### Biological Interpretation

The calculated probabilities reflect the likelihood that a given path of residues forms a connected route in a random subgraph of the protein contact network. Higher probabilities indicate paths that:

1. Traverse regions with high connectivity (residues with many contacts)
2. Have alternative parallel connections (increasing robustness)
3. Represent efficient communication routes through the protein structure

These properties are particularly relevant for understanding allosteric communication, where signal transduction often follows paths with high probability in the contact network [74].

### Computational Complexity

The most computationally expensive steps are:

1. Pseudoinverse calculation: O(n^3^) for n residues
2. Path enumeration: Exponential in worst case, but limited by path length constraint (3-7)
3. Determinant calculation: O(k^3^) for paths with k edges (k ≤ 6)

For typical proteins (n ≈ 100-500), the computation completes in seconds on standard hardware.

## Appendix E Signal-to-noise ratio with topology-weighted path probabilities

Given a protein represented as a graph G=(V,E) with vertices representing residues and edges representing physical contacts, the Burton-Pemantel theorem [39] provides the probability that a signal follows a specific simple path p between residues i and j. Let *w*_*p*_ denote this probability for path p, where:

w_p_=Prob(signal follows path p | ;signal travels from i to j)

The probabilities[39] {*w*_*p*_ } for all paths between i and j form a normalized distribution:

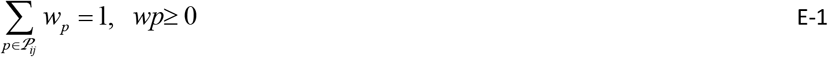

where 𝒫_*ij*_ is the set of ;all simple paths between residues i and j up to a maximum length L_max_.

### Signal and Noise Model

Consider a time-dependent signal S(t) originating at residue i. Each path p transmits the signal with probability wp (from the Burton-Pemantle theorem) and, when active, contributes independent thermal noise ηp(t) scaled by this probability.

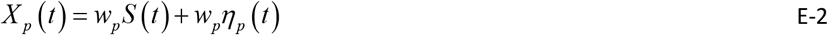

where ηp(t) are zero-mean Gaussian white noise processes with variance σ^2^, uncorrelated across different paths:

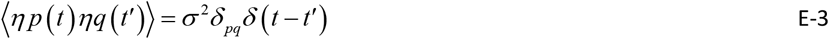

The received signal at residue j is the sum over all paths:

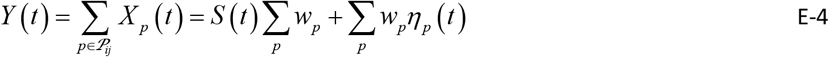

Since 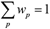 the signal component simplifies to S(t).

### Signal-to-Noise Ratio Derivation

The signal power is:

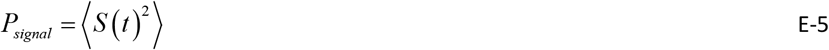

The noise power (variance) is:

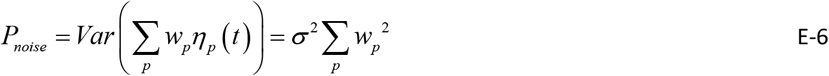

The signal-to-noise ratio (SNR) is therefore:

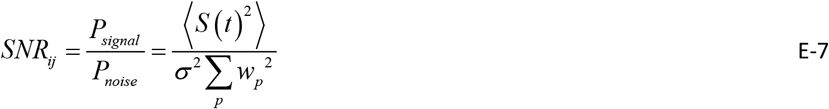

For comparative analysis between different residue pairs, we can consider the normalized SNR Where ⟨*S* (*t*)^2^ ⟩/ *σ* ^2^ = 1 :

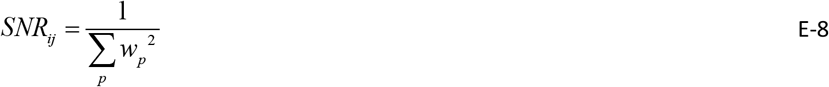

### Connection to Rényi Entropy

The denominator 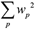 is precisely the collision probability of the path distribution, which is related to the Rényi entropy of order 2:

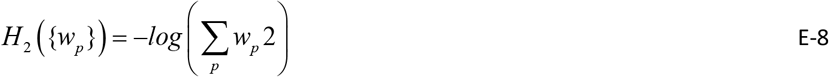

Therefore:

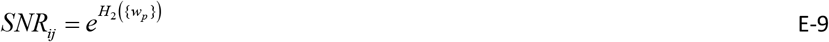

This shows that the SNR grows exponentially with the Rényi entropy of the path probability distribution. The Rényi entropy H_2_ provides an information-theoretic measure of the diversity in path utilization. Higher H_2_ indicates more uniform distribution of signal across paths, leading to better noise averaging.

